# Speed dependent descending control of innate freezing behavior in *Drosophila melanogaster*

**DOI:** 10.1101/234443

**Authors:** Ricardo Zacarias, Shigehiro Namiki, Gwyneth Card, Maria Luisa Vasconcelos, Marta A. Moita

## Abstract

The most fundamental choice an animal has to make when it detects a predator, or other threats, is whether to freeze, reducing its chances of being noticed, or to flee to safety. Here we show that *Drosophila melanogaster* exposed to looming stimuli in a confined arena either froze or fled. The probability of freezing versus fleeing was modulated by the fly’s walking speed at the time of threat, demonstrating that freeze/flee decisions were context dependent. We describe a pair of descending neurons crucially implicated in freezing. Genetic silencing of DNp09 descending neurons disrupted freezing yet did not prevent fleeing. Optogenetic activation of both DNp09 neurons induced running and freezing in a state-dependent manner. Our findings establish walking speed as a key factor in defensive response choices and reveal a pair of descending neurons as a critical component in the circuitry mediating selection and execution of freezing or fleeing behaviors.

## Introduction

Animals rely on similar kinds of cues to detect a predator’s rapid approach such as visual looming stimuli (Card, 2012; Carr, 2015; Pereira and Moita, 2016). Although great progress has been made in the study of threat detection mechanisms, much less is known regarding when and how different defensive behaviors are performed. Animals deploy several defensive strategies, grossly categorized into flight, freeze and fight, the display of which is modulated by external factors, such as the presence of offspring or existence of escape, and the animal’s internal state (Blanchard et al., 1986; Giaquinto and Volpato, 2001; Llaneza and Frye, 2009; Montgomerie and Weatherhead, 1988; de OCA et al., 2007a; Rickenbacher et al., 2017; Vale et al., 2017; Verma et al., 2016). How each specific defensive response is selected and executed remains unclear.

In mammals, mostly from studies using rodents, multiple brain regions have been implicated in the expression of freezing and escape responses including amygdala, hypothalamus and periaqueductal grey (Amorapanth et al., 2000; Gross and Canteras, 2012; Kim et al., 2013, 1993; Kunwar et al., 2015; Silva et al., 2013). More recently, microcircuits within these brain regions regulating the expression of flight or freeze behaviors have been characterized (Fadok et al., 2017; Tovote et al., 2016). Much less is known about the mechanisms of expression of defensive behaviors in other vertebrates. In fish, Mauthner cells, a pair of large neurons in the hindbrain, have been implicated in fast escape responses (review in (Korn and Faber, 2005), whereas other spinal chord projecting neurons are involved in slower escapes (Bhattacharyya et al., 2017). Even though the zebrafish habenula has been shown to down regulate freezing, favoring escape responses, how freezing behavior is produced remains unknown. In invertebrates the mechanisms of escape responses have received more attention. In striking similarity with the zebrafish, fruit flies can exhibit either fast or slower escape responses, the first relying on the giant fibers, a pair large descending neurons and the later on other descending neurons (Card, 2012; Fotowat et al., 2009; von Reyn et al., 2014). Although freezing behavior has been reported in fruit flies (Card and Dickinson, 2008b; Gibson et al., 2015), a systematic quantification of this behavior and exploration of its neural underpinnings is lacking.

While the neural circuits of defensive behavior have been partially described in multiple organisms, in mammals the focus has been on learned freezing responses, and in other organisms on innate escape responses. In addition, how external or internal factors impinge on these circuits to regulate choice between different responses remains largely unknown. To address this issue we decided to use *Drosophila melanogaster* for its arsenal of genetic tools to dissect neural circuits and the ability to use large sample sizes allowing for detailed quantitation of behavior. We developed a visual assay to track responses of flies to an expanding shadow mimicking a large object on a collision course - looming stimulus - that triggers defensive behaviors in virtually all visual animals tested, including fruit flies (Card, 2012; Carr, 2015; Pereira and Moita, 2016). Flies were exposed to multiple looming stimuli in an enclosed arena to increase the chances of seeing both escape and freezing responses. In our experimental set-up sustained freezing was the predominant defensive response. When flies did not freeze they displayed escape responses directed away from the looming stimulus. Taking advantage of a close-loop system, which allows the presentation of stimuli dependent on the behavior of flies, we found that the decision to freeze or flee was modulated by the flies’ walking speed at the time of threat. Through genetic manipulation of neuronal activity we identified a pair of descending neurons whose activity is required for freezing. Moreover, their ability to drive freezing, through optogenetic activation, depended on the walking speed of flies at the time of stimulation. These results reveal that innate responses to threats can be modulated by the flies’ internal state, while identifying an element of the freezing circuit that is modulated by this state.

## Results

### Flies jumped rarely in response to repeated inescapable looming

Figure 1 shows a schematic representation of the experimental setup. We placed single flies in a covered walking arena and gave them 5 minutes to explore. A computer monitor angled above the arena showed a looming stimulus (black circle expanding on a white background) repeated 20 times over a subsequent 5-min period. As a control, we showed a separate group of flies a sequence of randomly appearing black pixels resulting in a similar change in luminance but with no pattern of expansion (Figure 1B). Notably, flies cannot escape from the arenas.

**Figure 1.**
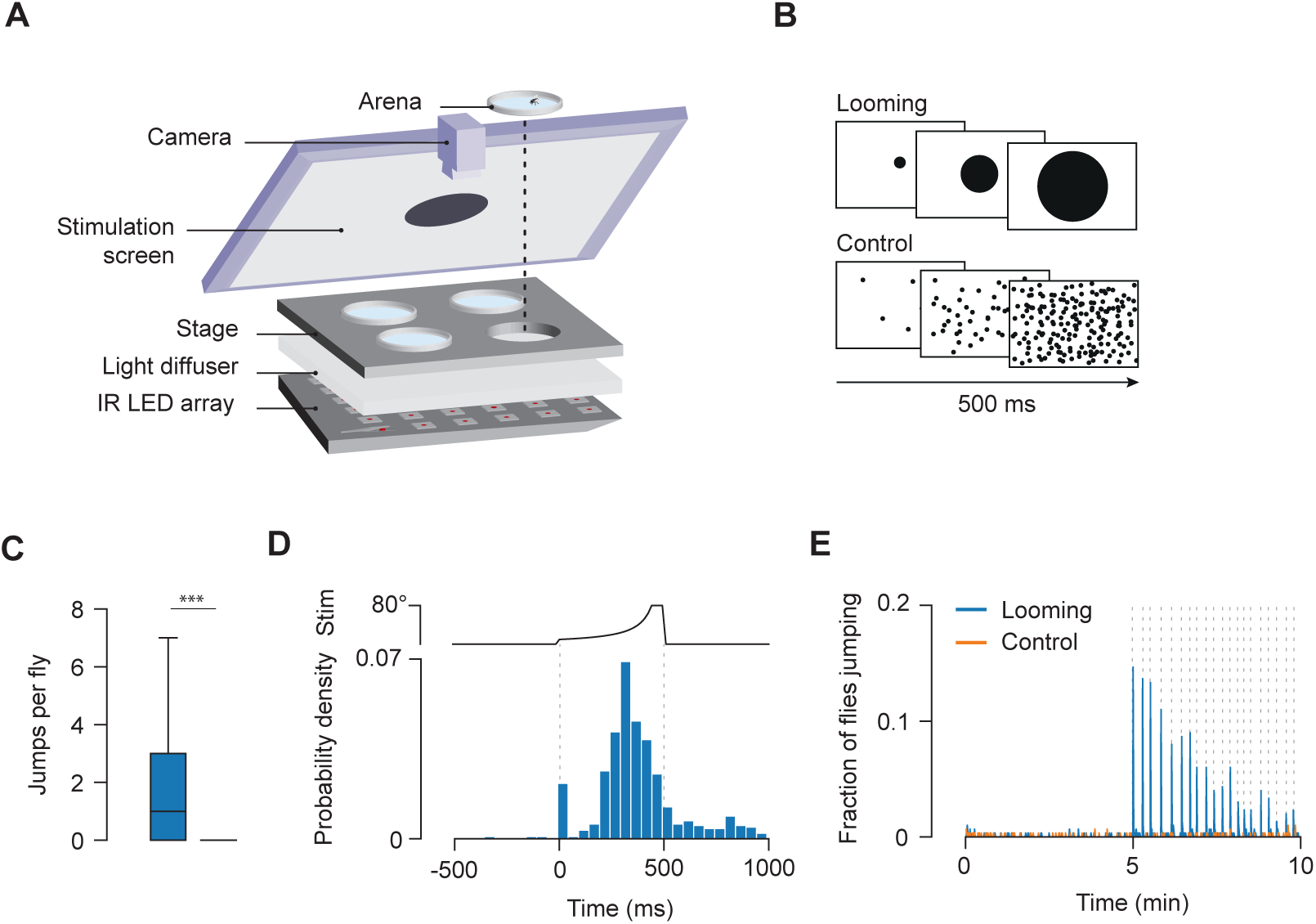
Flies jump in response to repeated looming. (A) Schematic of behavioral assay. (B) Schematic of visual stimuli, 300 flies tested in each condition. (C) Number of jumps per fly during stimulation period. *** denotes, p<0.0001. (D) Probability density of jumps around the looming stimulus. Dotted lines indicate beginning and end of looming. Top, visual angle of looming disk. (E) Fraction of flies jumping. Orange corresponds to flies exposed to control stimulus and blue to flies exposed to looming stimulus. Dotted lines represent stimulus presentations.

We found that looming stimuli only occasionally triggered escape jumps, or takeoffs (6.4% of looming stimuli, 384/6000), the most studied defensive response in insects (Card and Dickinson, 2008a, 2008b; Fotowat et al., 2009; Hammond and O’Shea, 2007; Heitler and Burrows, 1977; McKenna et al., 1989; von Reyn et al., 2014; Tauber and Camhi, 1995; de Vries and Clandinin, 2012). The number of jumps per fly was significantly higher for looming than control condition (Wilcoxon Rank Sum test Z= 14.15, p < 0.0001, Figure 1C) and the large majority of these events occurred within the window of stimulation, before the circle reached its maximum size (Figure 1D). Still, the efficacy of looming stimuli to elicit jumps was lower in our experimental conditions than that reported previously (Fotowat et al., 2009; de Vries and Clandinin, 2012), where flies were exposed to a single escapable looming stimulus. Furthermore, the probability of jumping decreased over the course of the 20 stimulus presentations (Figure 1E), suggesting that with multiple presentations flies may have habituated to looming. Alternatively, flies could be adopting other defensive strategies.

### Most flies responded to looming with sustained freezing

As running is an alternative form of defensive behavior (Lebestky et al., 2009), we analyzed the flies’ speed over the course of the experiment. The largest fraction of flies decreased their speed instead (Figure 2A). Visual inspection of the videos led to the observation that flies were not just walking slower, they were completely immobile, i.e. freezing, many times sustaining unbalanced postures for long periods of time (movies S1 and S2). In order to quantify freezing, we created an automated classifier based on pixel changes recorded in a region of interest surrounding the fly (Figure S1).

**Figure 2.**
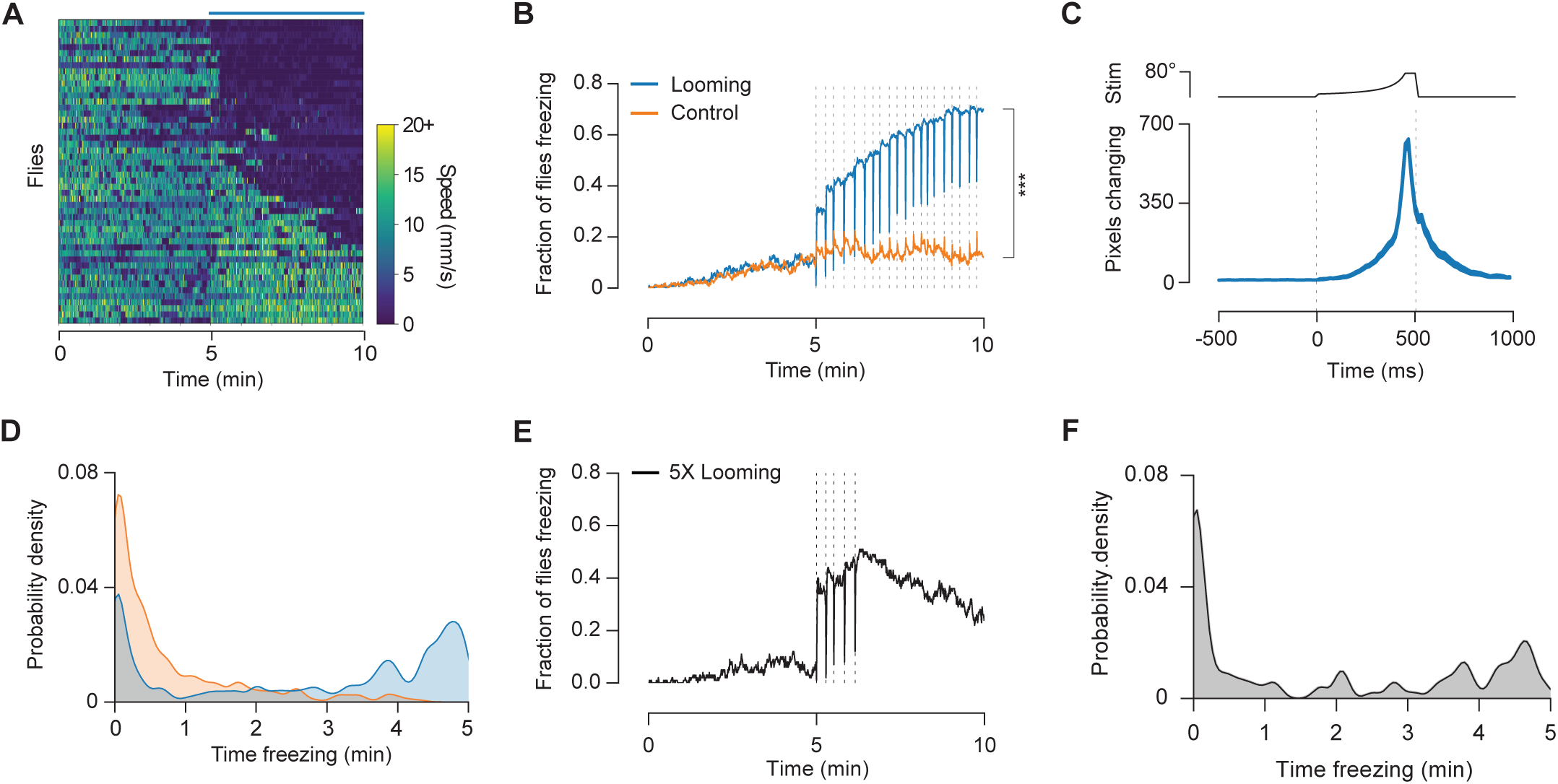
Flies freeze upon looming stimulation. (A) Speed in 500 ms bins for a subset of 50 flies. Each row corresponds to one fly, rank-ordered by time spent immobile. Blue bar on top indicates the 5 min stimulation period. In B-D orange corresponds to flies exposed to control stimulus and blue to flies exposed to looming stimulus. (B) Fraction of flies freezing. Dotted lines represent stimulus presentations. (C) Average (±sem) number of pixels changing around the fly, aligned on looming events where flies were freezing before and after the stimulus (n=1434). Dotted lines indicate beginning and end of looming. Top, visual angle of looming disk. (D) Probability density of time spent freezing by individual flies. (E) Fraction of flies freezing to a shorter stimulation (5 looming stimuli), n=100. Dotted lines represent looming presentations. (F) Probability density of percentage of time freezing during stimulation period for flies that received only 5 looming stimuli.

The fraction of flies freezing increased gradually with each looming presentation, arguing against habituation to the looming stimulus (Figure 2B). By the end of the experiment, 70% of flies were freezing compared to 12% for the control stimulus (Chi-squared test, X^2^ =208.599, p<0.0001, Figure 2B). Interestingly, there was a sharp decrease in freezing during stimulus presentation. Further examination showed that freezing flies displayed startle responses during looms, but quickly returned to an immobile state (movie S1). These responses were characterized by a spike in pixel motion that peaked at the end of the expansion of the looming stimulus (Figure 2C). Hence flies froze for long periods of time only breaking freezing for the brief moments of startle. Indeed, we observed a bimodal distribution, where most flies either froze for long periods of time (> 3 min) or did not freeze at all, with fewer flies displaying intermediate freezing levels (Figure 2D). To test whether prolonged freezing was due to the repeat stimulation, we exposed flies to five looming stimuli over the course of one minute, and asked whether flies would stop freezing after the stimulation period. We found that a substantial fraction of flies froze after the stimulation period (Figure 2E), and that 50% of the flies froze more than one minute with many flies freezing up to 5 minutes (Figure 2F). These long periods of time spent freezing are in sharp contrast with previous studies, which have only reported shortlived, occasional freezing in *Drosophila* (Card and Dickinson, 2008b; Gibson et al., 2015; Zabala et al., 2012).

### Flies that did not freeze, fled instead

Although a large fraction of flies froze in response to looming, not all flies did so. To determine whether flies displayed an alternate response to the looming stimuli, we next analyzed the flies’ behavior excluding all freezing and grooming bouts, hence only periods classified as walking (> 4 mm/s) were analyzed. During the baseline period walking speed gradually decreased, reflecting a common process of habituation to the test arena. However, during the stimulation period, their walking speed increased (one sample Wilcoxon Signed-Rank test, W=16435, p<0.0001). This was not observed for flies exposed to control stimuli, which further decreased their speed (one sample Wilcoxon Signed-Rank test, W=-15623, p<0.0001) (see Figure 3A and B). The average difference in walking speed between stimulation and baseline periods was significantly higher for flies exposed to looming relative to control (Wilcoxon Rank-Sum test, Z =16.33, p<0.0001 Figure 3B). In addition, we observed sharp increases in speed at the time of each looming stimulus (Figure 3A). To examine changes in speed around looming we plotted the average speed of walking flies aligned on looming onset (Figure 3C). Walking speed was relatively constant before the stimulus. Upon looming onset, flies initially paused for about 300ms followed by a rapid burst of locomotion (Figure 3C, marker #2, movie S3). These pauses may correspond to the brief freezing bouts preceding takeoff described previously (Card and Dickinson, 2008b; Zabala et al., 2012). Around the end of the looming stimulus a transient decrease in speed is seen. However, after the offset of the looming stimulus, the flies’ speed remained significantly elevated relative to the speed prior to looming (one sample Wilcoxon Signed-Rank test, W=208460, p<0.0001, Figure 3D). We next asked whether the running bouts reported here correspond to escapes directed away from the threat. We measured the orientation of the paths of walking flies before and after each looming presentation (Figure 3E). Before looming, paths in all orientations could be seen (Figure 3F). Upon looming, we observed a sharp increase in orientation bias toward the side of the chamber furthest away from the threatening stimulus (Figure 3G. Wilcoxon Signed-Rank test, W=178808, p<0.0001). Flies heading towards the screen changed direction, while flies heading away from the screen increased their walking speed and kept the same orientation. These findings indicate that running bouts triggered by looming stimuli, are not just a simple increase in locomotion, but constitute directed escape responses.

**Figure 3.**
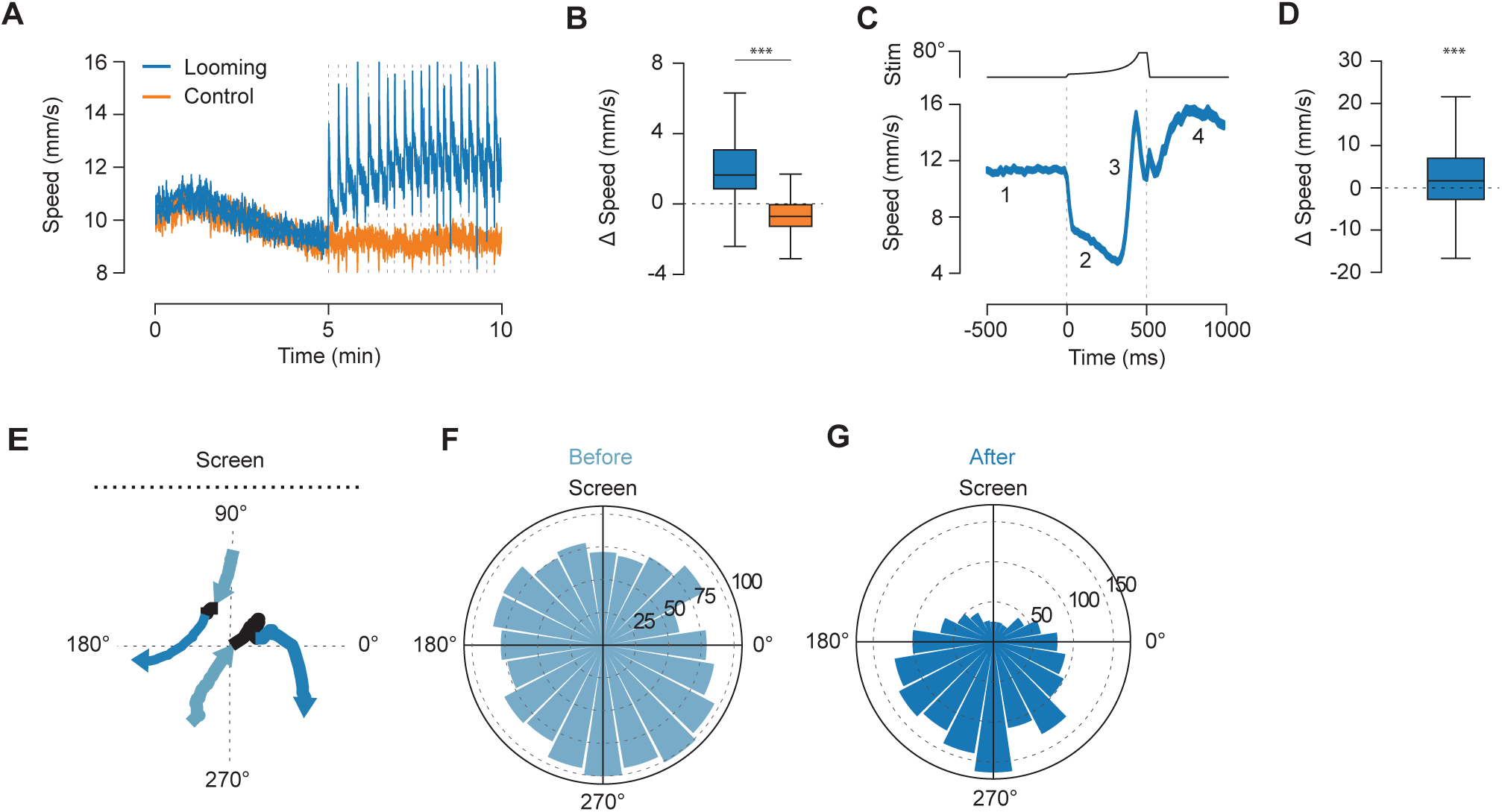
Flies run upon looming stimulation. A-G include walking bouts only (looming, n=280, and control, n=298). (A) Average walking speed (±sem). Dotted lines represent stimulus presentations. (B) Change in walking speed (baseline subtracted from stimulation period). In C-G n=1574 looming events. (C) Average speed (mean ± s.e.m.) around looming of walking flies. Numbers represent four stages of the response: 1-pre-looming, 2 - pause, 3 - run, 4 - post-looming. Top, visual angle of looming disk. (D) Change in walking speed (pre-stimulus baseline subtracted from post-stimulus period). (E) Two example paths during looming presentation within the chamber. Path before, during and after looming in light blue, black and dark blue, respectively. (F) Distribution of path orientation before looming. Bar heights indicate looming counts. (G) Distribution of path orientation after looming. *** denotes, p<0.0001.

### Freezing/fleeing decisions were modulated by the walking speed of the flies at the time of threat

Our data suggest that flies select between two distinct behavioral strategies, freezing or fleeing. A possible factor modulating this selection became apparent when we analyzed the relationship between the flies’ movement speed and their response to looming. We sorted looming trials by the speed of flies before looming onset and calculated the probability of freezing at different movement speeds. We observed a sharp decay in freezing probability with increasing speed, such that flies moving slowly or grooming were more likely to freeze upon looming stimulation than flies moving faster (Figure 4A). To further explore the relationship between movement speed at the time of looming and freezing probability, we designed a closed-loop experiment where the speed of the tested fly was tracked online and looming stimuli were delivered at specific speed thresholds. One group of flies received looming at low movement speeds and another at high movement speeds (<1mm/s and >15mm/s respectively. Figure S2). Importantly, there was no difference in the average baseline speed between the two groups, indicating that their overall level of arousal was similar (Student’s T-test, t=0.058, p=0.95, Figure 4B). We found that flies walking at low speed when exposed to looming were more likely to freeze than flies exposed to the same stimulus while walking faster (Wilcoxon Rank-Sum test, Z =-6.786, p<0.0001, Figure 4C), thus confirming a modulation of freezing probability by the flies’ movement speed. It is possible that the faster the flies walk the more difficult is to come to a fast halt, explaining the sharp decrease of freezing with increased speeds. However, we found that the probability of pausing upon looming was not significantly modulated by the flies walking speed (Figure S2). Indeed at all walking speeds, including the highest, flies paused in response to ~40% of the looming stimuli. Hence, the modulation of freezing probability is unlikely to result from a simple inability to become immobile at higher walking speed.

**Figure 4.**
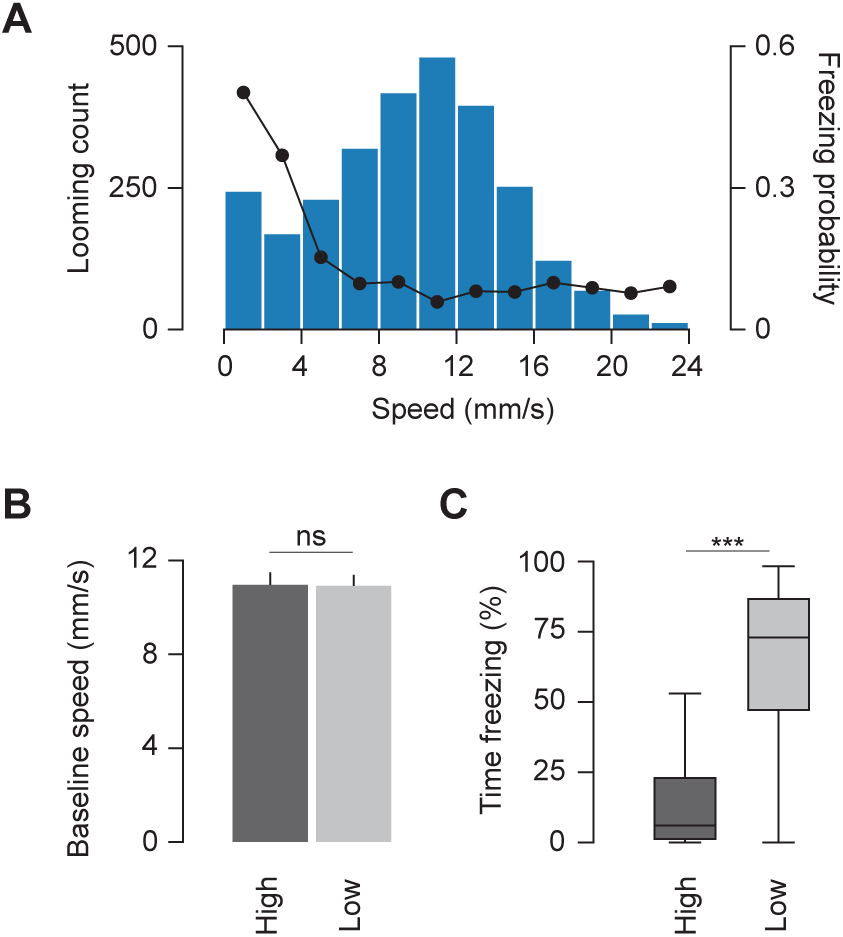
Response to looming is modulated by walking speed. (A) Distribution of speeds 500 ms before looming where flies were not freezing (blue bars, n=2740 loomings) overlaid with the probability of freezing after looming for each movement speed interval (black line with black dots). In a closed loop experiment flies were exposed to looming when moving slowly (low threshold, n=60) or fast (high threshold, n=56). (B) Average speed (mean ± s.e.m) during baseline period for high and low movement speed groups. (C) Percent time spent freezing during stimulation period for both groups. *** denotes, p<0.0001.

### Activity of DNp09 descending neurons is required for freezing but not fleeing in response to looming

Next, we searched for the neural mechanisms underlying the defensive behaviors observed. We focused on freezing, as it is conserved across the animal kingdom (Blanchard and Blanchard, 1969; Eilam, 2005a; Speedie and Gerlai, 2008) and corresponds to the dominant behavior adopted by flies in our experimental conditions. Given the lack of knowledge regarding the neural mechanisms of freezing in insects, we performed an unbiased screen, testing for looming triggered freezing of fly lines expressing a hyperpolarizing potassium channel, Kir2.1 (Baines et al., 2001), in different subsets of descending neurons (DNs; Namiki et al, in preparation, <u>Janelia Descending Interneuron Project</u>). We focused on DNs, as these convey information from the brain to the ventral nerve cord thereby controlling behavior. From this screen we identified a bilateral pair of DNs, DNp09, with dendrites innervating the posterior protocerebrum and a large axon extending throughout the ventral nerve cord, as well as the posterior slope and gnathal ganglion (Figure 5A. Weak off-target labeling not shown). Silencing these neurons significantly decreased the occurrence of freezing relative to the parental controls (Kruskal-Wallis test, X^2^=80.19, p<0.0001. Post-hoc Dunn tests revealed a significant difference between DNp09>Kir2.1 and the parental controls: DNp09>Kir2.1 vs. DNp09/+, Z=-4.98, p<0.0001; DNp09>Kir2.1 vs. UASKir2.1/+, Z=-8.94, p<0.0001. In addition the two parental controls showed a small but significant difference: DNp09/+ vs UASKir2.1, Z=3.96, p=0.0002. Figure 5B and C). This effect was not due to an overall decrease in sensitivity to looming since running was intact (Figure 5D and Figure S4A). Furthermore, the frequency of jumps was increased, raising the possibility that in control flies freezing behavior directly or indirectly inhibits jumps (Figure S4B).

**Figure 5.**
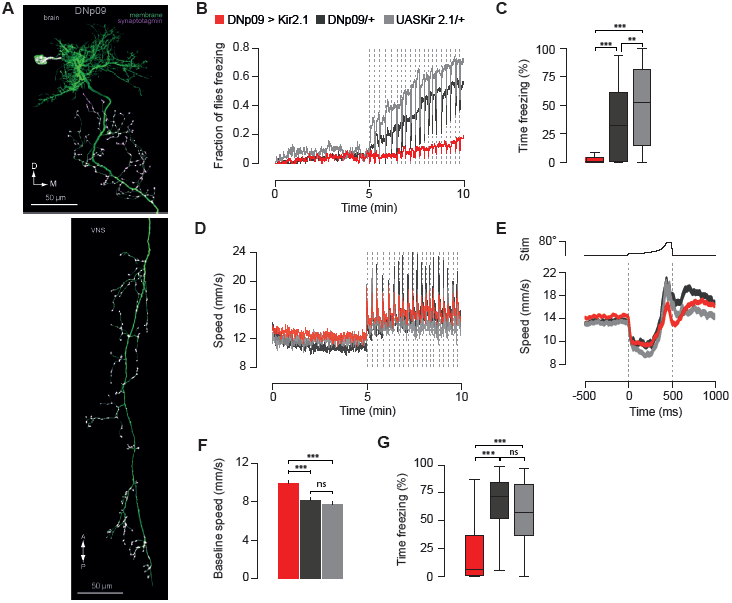
Silencing DNp09 descending neurons affects freezing but not running. (A) We used a split-GAL4 driver line specific for DNp09 neurons (SS1540). DNp09 morphology is depicted in this confocal image of the driver line crossed to a GFP marker with anti-synaptotagmin-HA staining. B-F include n=100 flies per genotype. (B) Fraction of flies freezing for flies with DNp09 neurons silenced (red line) or not manipulated (gray lines correspond to parental controls). Dotted lines represent stimulus presentations. (C) Percent time spent freezing during stimulation period for each genotype. D-E shows speed for periods classified as walking for flies with DNp09 neurons silenced. (D) Average walking speed (±sem). Dotted lines represent stimulus presentations. (E) Average speed (mean ± s.e.m.) around looming. Dotted lines indicate beginning and end of stimulus. (F) Average baseline speed (mean ± s.e.m.) for each genotype. (G) Percent time spent freezing in closed-loop when looming stimulation was restricted to low speeds. DNp09>Kir2.1, n=42; DNp09/+, n=49; UASKir2.1/+, n=50. ** denotes p<0.001 and *** p<0.0001.

### DNp09-silenced flies still paused in response to looming

The effect of silencing DNp09 neurons on freezing behavior could reflect a general impairment in the ability to stop. However, DNp09-silenced flies that ran in response to looming still exhibited a pause upon looming onset, which can be seen in the walking speed profiles around looming stimulation (Figure 5E). Indeed, these flies paused in 30% of the looming stimuli (looming stimuli that occurred while flies were already freezing were excluded from this analysis). This finding indicates that pausing and sustained freezing are mediated by different mechanisms. Interestingly this dissociation is also present in rodents that possess distinct descending neurons for freezing and stopping, and where freezing involves sustained muscle tension whereas stopping does not (Bouvier et al., 2015; Koutsikou et al., 2014; Tovote et al., 2016).

### Disruption of freezing was independent of walking speed

Given the negative relationship between speed and freezing probability mentioned above, we investigated whether silencing DNp09 neurons affected the walking speed of the flies. We found that indeed the average baseline speed was increased in silenced flies relative to controls (One-way ANOVA, F=16.37, p<0.0001. Post-hoc Tukey revealed a significant difference between DNp09>Kir2.1 and the parental controls: DNp09>Kir2.1 vs. DNp09/+, p<0.0001; DNp09>Kir2.1 vs. UASKir2.1/+, p<0.0001. No difference between parental controls was found, p=0.6. Figure 5F). This raises the possibility that the impairment seen in freezing results from the increased walking speed and hence a shift in the probability of freezing behavior. Therefore, we calculated the probability of freezing for looming stimuli occurring at different speeds. We found that despite the upward shift in speed of DNp09-silenced flies relative to controls, freezing probability of these flies was lower for the entire range of speeds (Figure S4C). In addition, we tested DNp09-silenced and control flies in closed loop such that looming stimuli were only presented when flies were at very low speeds (1mm/s) and found that even at these speeds DNp09-silenced flies froze less than controls (Kruskal-Wallis test, X^2^=33.55, p<0.0001. Post-hoc Dunn tests revealed a significant difference between DNp09>Kir2.1 and the parental controls: DNp09>Kir2.1 vs. DNp09/+, Z=-5.64, p<0.0001; DNp09>Kir2.1 vs. UASKir2.1/+, Z=-4.16, p<0.0001. No difference between parental controls was found, p=0.36. Figure 5G). Together these findings indicate that silencing DNp09 neurons directly disrupts freezing, rather than indirectly affecting freezing behavior by increasing the speed of locomotion.

### Activation of DNp09 neurons induced freezing

If indeed DNp09 neurons are involved in the execution of freezing behavior, activating them artificially, in the absence of looming stimuli, should induce freezing. We expressed the red-shifted channelrhodopsin, CsChrimson (Klapoetke et al., 2014), in DNp09 neurons and exposed single flies to red light using a modified version of our behavioral setup (Figure 6A). CsChrimson requires retinal to function, thus experimental flies were raised in food containing retinal whereas control animals were raised in standard food. Flies were allowed to acclimate to the arena for 2 minutes. We then presented 10 trials of continuous light for 2 seconds, separated by 20-second intervals. We found that CsChrimson activation of DNp09 neurons was sufficient to trigger freezing in ~ 60% of trials (Figure 6B and movie S4). Closer inspection of the time course of freezing induction showed that the probability of freezing increased gradually over the course of the 2-second light stimulation (Figure 6C). The lag in freezing correlated with an initial increase in walking speed, induced by DNp09 activation (Figure 6D). This running bout was much reduced in control flies, showing that running was mostly caused by DNp09 activity (Figure S5). In an unbiased behavioral representation of optogenetic activation of descending neurons it was also observed that activation of DNp09 induced running followed by pausing using different analytical methods (Cande et al., 2017). The finding that flies first ran in response to DNp09 activation contrasts with the observation that, upon looming, flies often first paused and then ran, jumped or froze. Given that looming triggered pauses were independent of DNp09 neurons, it is possible that, with looming, neurons upstream of DNp09 inhibit the initial running bout seen with artificial DNp09 activation. Finally, we observed jumps at light offset in test flies, but not control flies (Chi-squared test, X^2^=219.76, p<0.0001, Figure 6E). Moreover, jumps were more likely after stimulations that led to freezing than after stimulations that failed to elicit freezing (Chi-squared test, X^2^=72.51, p<0.0001, Figure 6E). One possible explanation for this could be that strong activation of DNp09 neurons inhibits downstream targets involved in jumping behavior, such that when DNp09 activation stops, these neurons are released from inhibition showing rebound excitation, thereby triggering the observed jumps.

**Figure 6.**
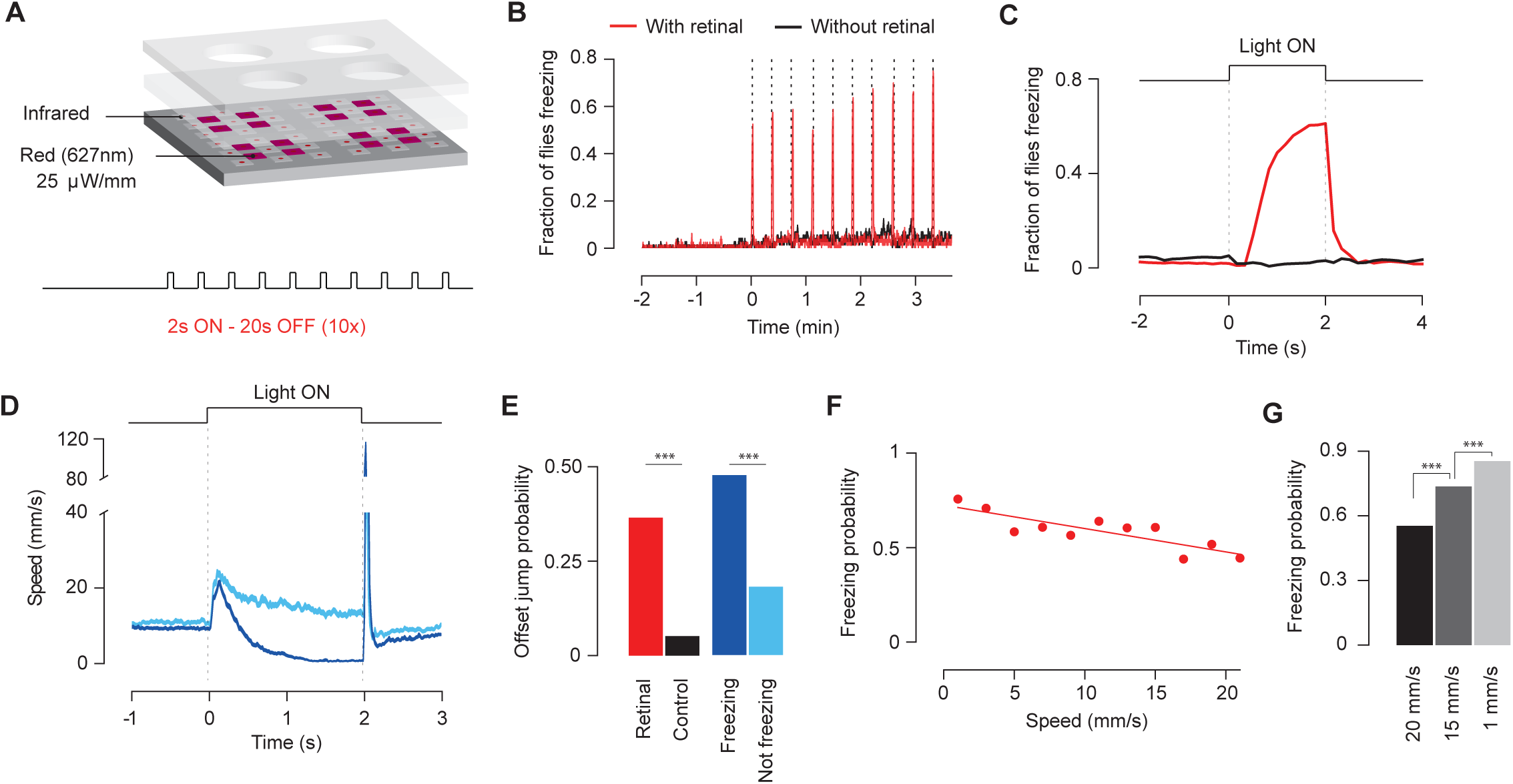
Activation of DNp09 descending neurons leads to freezing. (A) Schematic of experimental set-up for chrimson stimulation and stimulation protocol. Test flies supplemented with retinal (n=80) and control flies without (n=72). (B) Fraction of flies freezing. Vertical doted lines indicate light stimulation. (C) Fraction of flies freezing aligned on light activation. In e-g n=800 stimulation events with retinal, and n=720 stimulation events without. (D) Average speed (mean ± s.e.m.) aligned on light presentation for events that induced freezing (dark blue) and events that did not (light blue). (E) Probability of jumping at light offset for control and test flies (black and red bars). Probability of jumping at light offset for stimulation events of test flies that induced freezing and events that did not (dark, n=492, and light blue, n=308, bars). (F) Linear regression of probability of freezing by test flies upon red light stimulation at different pre-looming speeds. (G) Probability of freezing for DNp09>CsChrimson tested at different speeds (number of stimulation events for very high, high and low n=403, 674 and 593, respectively). *** denotes p<0.0001

### The probability of freezing upon DNp09 activation was dependent on the walking speed of flies

Given that the probability of freezing in response to looming stimuli was found to depend on the walking speed of flies at the time of threat, we asked whether the ability of DNp09 neurons in driving freezing was also modulated by the flies’ walking speed. We found that the probability of freezing upon light activation of DNp09 neurons was negatively correlated with the movement speed of flies (Figure 6F, linear regression, r^2^=0.71 p=0.001). To confirm that DNp09-driven freezing is modulated by the movement speed of flies at the time of stimulation, we again tested flies in close loop, such that DNp09 activation would occur when flies were at low, high or very high speeds (1mm/s, 15mm/s and 20mm/s, respectively). The probability of DNp09 activation driving freezing was highest for the flies stimulated at low movement speed (very high vs. high: Chi-squared test, X^2^ =37.875, p<0.0001; high vs. low: Chi-squared test, X^2^ =26.278, p<0.0001. p values Bonferroni corrected. Figure 6G). Together, these findings show that DNp09 neurons are a key element in the circuit mediating the speed modulation of freezing expression, and suggest that this modulation is not a result of locomotion induced changes in visual perception (Chiappe et al., 2010; Maimon et al., 2010).

## Discussion

Freezing is a defensive response characterized by complete immobility, allowing preys to avoid being detected while remaining attentive to changes in the environment (Eilam, 2005b; Fanselow, 1994). This response has been reported across vertebrates (Blanchard and Blanchard, 1969; Eilam, 2005b; Speedie and Gerlai, 2008). Here we show long lasting freezing in fruit flies similar to that observed in vertebrates and distinct from the brief freezing periods reported for flies (Card and Dickinson, 2008b; Gibson et al., 2015; von Reyn et al., 2014). The pervasiveness of freezing across distant taxa strongly suggests its independent evolution and supports its adaptive value (Schultz and Endler, 1987).

Given the prevalence of freezing behavior, it is crucially important to understand whether there are general principles that govern it. For instance, the display of freezing is plastic. Rodents tend to freeze only if there is no escape, and the presence of conspecifics decreases this behavior (Blanchard et al., 1986; Kiyokawa et al., 2007, 2009; de OCA et al., 2007b; Rickenbacher et al., 2017; Vale et al., 2017). Furthermore, it has been previously shown that estrous cycle or feeding state modulate freezing (Llaneza and Frye, 2009; Verma et al., 2016). How animal’s surroundings or the animal’s internal state regulate this behavior is much less clear. A hint of the conserved nature of the principles governing freezing, is the finding that in other vertebrates, such as fish, the expression of innate defensive behaviors is also plastic (Agetsuma et al., 2010; Bass and Gerlai, 2008; Giaquinto and Volpato, 2001). In this study, we extend this to invertebrate animals by demonstrating the plastic nature of freezing in flies. Flies either ran or froze in response to inescapable looming.

The choice between escaping and freezing was strongly modulated by the flies’ speed at the time of threat. This effect could be explained by an impact of speed on motor output, sensory processing and/or arousal. An effect on motor output could be simply a consequence of increased difficulty in stopping when walking fast. The findings that pausing in response to looming was independent of the walking speed, and that DNp09 induced freezing was always preceded by running, argues against this possibility. This leaves a possible influence on visual processing (Bennett et al., 2013; Chiappe et al., 2010; Maimon et al., 2010; Vinck et al., 2015) or central motor commands. The finding that DNp09-induced freezing was modulated by movement speed of flies at the time of stimulation argues for the later. Future experiments are required to disambiguate between these scenarios.

We next explored the neuronal underpinnings of freezing behavior, contributing to the understanding of how different animals, with different bodies and brains, implement this seemingly simple behavior. We uncovered the key role of a single pair of descending neurons, DNp09, in driving freezing. DNp09 neurons innervate visual input areas in the central brain, thus being in a good position to respond to looming stimuli. The output terminals of DNp09 neurons innervate the posterior slope, which is densely innervated by other descending neurons (Hsu and Bhandawat, 2016), and multiple regions along the ventral nerve chord, allowing the interaction with other motor outputs at different levels.

Although freezing is often seen as absence of other behaviors, a passive state of immobility (Fadok et al., 2017; Fanselow, 1994; Gozzi et al., 2010), evidence suggests otherwise. For example in mammals, freezing is accompanied by sustained muscle tension likely involved in postural control (Koutsikou et al., 2014; Watson et al., 2016) and, in response to learned cues, requires sustained activity of several brain regions (Amano et al., 2011; Burgos-Robles et al., 2009; Duvarci et al., 2011). In addition a recent study identified in mice a set of descending neurons that drive stopping behavior that is distinct from those identified for freezing (Bouvier et al., 2015; Tovote et al., 2016). The finding that looming triggered-freezing and pausing could be dissociated supports the idea that freezing is an active defense module pointing to the conserved nature of the distinction between freezing and stopping. Moreover, freezing may require active inhibition of alternate behaviors. An indication that active inhibition of alternate behavior happens in flies comes from our observation that DNp09 silenced flies jump more and that flies jump at the offset of DNp09 neuron activation, presumably resulting from rebound excitation after inhibition of jump. Consistent with an inhibitory effect of freezing on jumping is the observation that jumping steeply decreased with increase number of flies freezing over the course of the repeated looming stimuli. The identification of DNp09 descending neurons as central to freezing opens the path to further explore how the active state of freezing is implemented. Activation of DNp09 neurons drove both running and freezing. However, silencing DNp09 neurons left looming triggered escape responses intact suggesting that DNp09 triggered running may correspond to a different behavior. Still, it will be very interesting to unravel how a single pair of neurons drives distinct behaviors.

Finally, since the flies’ speed modulates their response to looming stimuli, we examined whether the ability of DNp09 neurons to drive freezing was also modulated by the flies’ speed. We found that the probability of freezing upon DNp09 stimulation was negatively correlated with the flies’ movement speed. This finding demonstrated that DNp09 neurons are a key element in the circuit mediating speed dependent defensive decisions. Unraveling how speed impinges on DNp09 neurons and possibly other elements of defense circuits will be instrumental for the understanding of the organization of defensive behaviors crucial for survival.

## Acknowledgements

We would like to thank the Scientific Software Platform of Champalimaud Centre for the Unknown (CCU) for developing the fly tracking software; the Scientific Hardware Platform of the CCU, for building LED array for optogenetic activation experiments; the Fly facilities of the CCU and Grace Zheng from the fly facility of Janelia Research Campus; Eugenia Chiappe and Joseph Paton for comments on the manuscript. This work was funded by the Champalimaud Foundation, the visiting scientist program of Janelia Research Campus and the ERC Starting Grant CoCO 337747.

## Author Contributions

RZ performed all experiments and analyzed data. RZ, MLV and MAM designed all experiments, discussed results and wrote manuscript. SN and GC created the split-gal4 lines of descending neurons and provided the image of DNp09 neuron labeling. GC, MLV and MAM supervised the unbiased DN-silencing screen performed at Janelia Research Campus; GC commented on the manuscript. Fly lines should be requested to GC. All data are reported in the main text and supplementary materials, stored at Champalimaud Research and available upon reasonable request.

## Experimental Procedures

### Animal husbandry and fly strains

All animals used in experiments were 4-6 day old mated female *Drosophila melanogaster*. Flies were raised at 25°C and 70% humidity in a 12h:12h dark:light cycle. Strains and sources:Canton S used in in figures 1, 2, 3 and 4; DL used to cross with parental strains in figure 5 and 6: 10XUAS-IVS-eGFPKir2.1 (attP2) (DL) (DL denotes chromosomes derived from the DL fly strain)(von Reyn et al., 2014); 20xUAS-CsChrimson.mVenus in attp2(Klapoetke et al., 2014); DNp09 line (Namiki et al, in preparation; https://www.janelia.org/project-team/fly-descending-interneuron).

### Behavior

All behavioral experiments were performed in the 4-hour period preceding lights off and under the same conditions as rearing.

### Arenas

Behavioral arenas were custom built from opaque white and transparent acrylic sheets. Chambers were 30 mm in diameter and 4 mm in height.

### Apparatus

We recorded behavior of unrestrained flies while presenting visual stimulation (Figure 1a). A monitor tilted at 45 degrees over the stage delivered visual stimulation. To image fly locomotion, a custom-built infrared (850 nm) LED array was placed under the stage (backlight). A diffuser (2 mm white opaque acrylic sheet) was placed on top of the LED array to produce homogeneous illumination. Fly behavior was recorded using a USB3 camera (PointGrey Flea3) with a 850 nm long pass filter. Each fly was aspirated into a chamber and placed on the stage. Flies were observed for 20 seconds to ensure that no gross motor defects were present before video acquisition started.

### Video acquisition and tracking

Videos were acquired using Bonsai(Lopes et al., 2015) at 60 Hz and width 1104 x height 1040 resolution. Image segmentation was performed by custom software in python using OpenCV. Fly positions were calculated from centroid, and motion quantified by pixel change in an 100x100px region of interest surrounding the centroid.

### Visual stimulation

Visual stimuli were presented on a 24-inch monitor running at 144 Hz (ASUS VG248QE). All stimuli were generated in a custom python script using PsychoPy(Peirce, 2007).

Looming: The visual angle of the expanding circle was determined by the equation θ (t) = 2 tan-^1^(l/vt)

where l is half of the length of the object and v the speed of the object towards the fly. Virtual object length was 1 cm and speed 25 cm/s (l/v value of 40 ms). Each looming presentation lasted for 500 ms. Object expanded during 450 ms until it reached maximum size of 78 degrees where it stayed for 50 ms before disappearing.

Random dots: an array of approximately 3 degree dots was added each frame in random positions to generate similar change in luminance as the looming but without any expanding pattern.

### Behavioral classifiers

Jumping: a fly was classified as having jumped if frame by frame speed exceeded a threshold (75 mm/s) identified by a discontinuity in the speed distribution.

Freezing: a fly was considered to be freezing when average pixel change in a region of interest around the fly was lower than 25 pixels (approx.< 5% of fly area) over 500 ms.

Walking: a fly was considered to be walking if its average speed over a 500 ms period exceeded 4 mm/s.

### Closed-loop looming stimulation

Fly positions were tracked in real time and used to trigger looming stimuli using Bonsai(Lopes et al., 2015). Threshold for looming stimuli were defined based on the path length in 500 ms windows. Low speed loomings were triggered when path length was < 1 mm in 500 ms. High speed loomings were triggered when path length was > 7.5 mm in 500 ms. A refractory period of 15 seconds was imposed after each triggered stimulation.

### Optogenetic activation

Responses of freely moving flies to CsChrimson(Klapoetke et al., 2014) activation were captured using a modified version the behavioral apparatus described in figure 6a. High-powered 627 nm LEDs were interspersed between the infrared LEDs on the backlight board. Each arena was irradiated with four LEDs for total radiance of 0.025mW.mm^−2^. Experimental flies were raised on standard fly food with 0.2 mM all trans-retinal (Sigma, R2500) and control flies were raised on standard fly food without retinal. Flies were allowed to explore for 2 minutes and then were stimulated with 10 repetitions of 2 seconds light on, 20 seconds light off.

### Closed-loop optogenetic activation

Fly positions were tracked in real time and used to trigger red light activation using Bonsai(Lopes et al., 2015). Speed thresholds used were the same as in the looming stimulation. Threshold for looming stimuli were defined based on the path length in 500 ms windows. Low speed loomings were triggered when path length was < 1 mm in 500 ms. High speed loomings were triggered when path length was > 7.5 mm in 500 ms. A refractory period of 15 seconds was imposed after each triggered stimulation.

### Staining and imaging

Imaging of DN morphology was performed as part of the Janelia Descending Interneuron project. The DNp09DN split-GAL4 driver line, SS1540, was crossed to 5XUAS-IVS-Syt∷smGFP-HA and-5xUAS-IVS-myr∷smGFP-FLAG and the central nervous systems of the progeny were dissected and stained for anti-GFP according to the standard Janelia FlyLight protocol. Brains were subsequently mounted in DPX and imaged with a confocal microscope. Detailed immunohistochemistry staining and DPX mounting protocols are avilable online at https://www.janelia.org/project-team/flylight/protocols. To best illustrate DNp09 morphology, off-target expression was removed from the image using Photoshop.

### Data analysis and statistics

Data analysis was performed using custom Python scripts. All data, except those from animals excluded due to tracking errors, were analyzed. Prior to statistical testing, data were tested for normality with a Shapiro-Wilk test and the appropriate non-parametric test was chosen if data were not normally distributed. All statistical tests are specified in the results section of the text or figure captions and are two-sided.

